# Hydrodynamic dissection of single cells in a microfluidic system

**DOI:** 10.1101/2022.06.09.495579

**Authors:** Rajorshi Paul, Kevin S. Zhang, Myra Kurosu Jalil, Nicolas Castaño, Sungu Kim, Sindy K.Y. Tang

## Abstract

*Stentor coeruleus*, a single-cell ciliated protozoan, is a model organism for wound healing and regeneration studies. Despite *Stentor*’s large size (up to 2 mm in extended state), microdissection of *Stentor* remains challenging. In this work, we describe a hydrodynamic cell splitter, consisting of a cross junction, capable of splitting *Stentor* cells in a non-contact manner at a high throughput of ∼500 cells/min under continuous operation. Introduction of asymmetry in the flow field at the cross junction leads to asymmetric splitting of the cells to generate cell fragments as small as ∼8.5 times the original cell size. Characterization of cell fragment viability shows reduced 5-day survival as fragment size decreases and as the extent of hydrodynamic stress imposed on the fragments increases. Our results suggest that cell fragment size and composition, as well as mechanical stress, play important roles in the long-term repair of *Stentor* cells and warrant further investigations. Nevertheless, the hydrodynamic splitter can be useful for studying phenomena immediately after cell splitting, such as the closure of wounds in the plasma membrane which occurs on the order of 100 – 1000 seconds in *Stentor*.

## 1. INTRODUCTION

Wound healing and regeneration are vital biological processes essential for homeostasis and ultimately, for survival. They occur in biological systems of widely varying length scales, from the single cell level^1-9^ to the tissue level.^5, 10-13^ The mechanisms of wound repair differ in different types of cells or tissues.^6, 8, 13, 14^ While the importance of wound healing at the single cell level is gaining recognition, the precise mechanisms involved are not yet fully understood.^8^ *Stentor coeruleus*, a giant single-cell ciliated protozoan (size ranging from 200 µm to 400 µm in contracted state, and up to 2 mm in extended state), shows remarkable wound healing and regeneration capabilities.^7-9, 15-21^ *Stentor* has been shown to survive wounds as large as 100 μm in diameter that span nearly half the size of the cell.^7^ It exhibits one of the highest wound healing rates (8 = 80 μm^2^/s) among previously studied single cell models.^1, 4, 7, 9, 14, 22-25^ *Stentor* has also been observed to employ unique, large-scale, mechanical behaviors that may facilitate its wound healing process.^7^ In addition to their ability to heal large wounds, *Stentor* cells are capable of regenerating into fully functioning cells from fragments as small as 1/27^th^ the size of the original cell.^16^ The highly polyploid macronucleus in *Stentor coeruleus* is thought to be a contributing factor to its robust regenerative capability.^17^ This trait may allow even a small fragment of the cell to carry sufficient copies of the genome to enable regeneration.^8, 18, 19, 21, 26, 27^ However, the detailed mechanisms of wound healing and regeneration in *Stentor* remain incompletely understood. One of the major experimental challenges is the lack of a high-throughput and repeatable method to wound and/or to dissect the cells.

Microdissection of small organisms has traditionally been performed manually since its inception in the 1600s.^28, 29^ From the late 1800s to today, microdissection of single cells has primarily utilized manually controlled microneedles for cutting while the process is monitored under an optical microscope.^30-32^ Manual surgery using microneedles poses a few challenges. First, it is imprecise and does not allow cell alignment or reproducibility of cuts along cell axes. Second, manual surgery is a slow process. Depending on the skill of the operator, cutting each cell can take as long as 3 minutes.^9^ Analyses such as proteomics and RNA sequencing often require hundreds of cells. Manual dissection of hundreds of cells would require many hours, which is much longer than the time scale in which *Stentor* cells heal from wounds and begin to regenerate from fragments. This dissection rate limits the ability of downstream analysis to capture transient biological processes, e.g., transcription and translation events, related to wound healing and regeneration.

Microfluidics offers a way to perform single cell dissection in a repeatable manner with increased throughput. Recently, our group has developed a microfluidic guillotine that bisects single cells in a flow-through manner using a solid poly(dimethylsiloxane) (PDMS) blade inside a microfluidic device.^7, 9^ The microfluidic guillotine is nearly 200 times faster than manual surgery with a cutting time of approximately 1 second for each *Stentor* cell. The device can generate reproducible wound patterns while preserving cell viability post-wounding (up to 95% cell viability 1 day following bisection). However, debris build-up at the tip of the PDMS blade can prevent long-term use of the microfluidic guillotine, limiting the number of cells that can be bisected per experiment to less than 20.^9^

In this paper, we describe the design, development, and characterization of a hydrodynamic cell splitter capable of high-throughput splitting of *Stentor* cells without debris build-up. The device consists of a cross junction where cells are split by hydrodynamic extensional stresses. The device splits *Stentor* cells at a throughput of approximately 500 cells per minute, which is over 8 times faster than the microfluidic guillotine and over 1500 times faster than manual surgery. As the device relies only on hydrodynamic forces to split cells without using any solid surfaces, this design completely avoids clogging from cellular debris, and the device can be operated continuously without fouling. Introduction of asymmetry in the flow field at the cross junction causes asymmetric splitting of the cells. We also characterize the degree of wounding and 5-day survival of the cell fragments generated by both the splitter and the guillotine. Our investigation indicates that hydrodynamic stresses adversely affect the long-term cell viability post-splitting and may contribute to unusual cellular behavior.

## 2. EXPERIMENTAL DESIGN

### *Stentor* and algae cultures

*Stentor coeruleus* culture was obtained from Carolina Biological Supply and was prepared by modifying the standard protocol for *Stentor* cultures.^32^ Cultures were grown in the dark at 22 °C in pasteurized spring water (PSW) (132458, Carolina Biological Supplies) in clean 400 mL Pyrex dishes. *Stentor* were fed algae *Chlamydomonas reinhardtii* every two days (∼0.2 μL of algae per *Stentor* cell in the culture). The *Stentor* cultures were cleaned approximately on a monthly basis to prevent accumulation of waste materials in the culture. The culture medium was replaced with fresh PSW every two months. A liquid culture of *C. reinhardtii* was grown separately under constant room light in standard Tris acetate phosphate (TAP) medium and washed twice with PSW before being fed to *Stentor*. The liquid algae cultures were replaced every 4-5 days when the cells became barely motile.

### Microfluidic device fabrication and operation

The detailed schematic diagram and dimensions of the hydrodynamic splitter are shown in Fig. S1. The microfluidic guillotine used in this work was the same as that reported previously.^7^ The microfluidic devices were designed in AutoCAD 2020 and fabricated using standard techniques of soft lithography. The heights of the master molds were measured with a profilometer to be 100±8 µm. The final hydrodynamic splitter devices were made of two layers of PDMS (SYLGARD-184, Dow Corning) cured from the master molds (10:1 base-to-curing agent) and bonded to a glass slide using a plasma cleaner. The devices were heat treated at 120 °C for 15 min immediately after bonding to strengthen the adhesion between the PDMS layers, and between the PDMS layer and the glass slide.

Before running the experiments, the devices were primed by first washing them with ethanol and then with PSW. Next, they were placed in a Pyrex glass dish containing enough PSW to submerge the microfluidic device. The dish was placed inside a desiccator under vacuum for 8 hours to remove any remaining air bubbles by gas permeation through PDMS. The single-layered microfluidic guillotine did not require this additional desiccation step. In between experiments, the channels were washed of any debris with PSW and discarded if the channels remained clogged by debris.

*Stentor* cells used in the experiments were pipetted into 4 mL glass vials from the culture. To prevent cells from getting wounded, the narrow ends of the pipette tips were cut off to ensure that the diameter of the tip was larger than the dimensions of the cells. The cells in the glass vials were then carefully transferred into polyethylene tubing (BB31695-PE/4, Scientific Commodities Inc., inner diameter = 762 μm) attached to a 12 mL Monoject plastic syringe filled with PSW. A syringe pump was used to inject the cells into the microfluidic device at the desired flow rates. To ensure that the flow was steady inside the device before the cells entered, the cells were loaded into the tubing as far from the device as possible such that a sufficient amount of PSW (>100 μL) would flow into the device before the cells entered. For the hydrodynamic splitter, the flow rate was varied between 25 mL/h and 300 mL/h at the inlet, which was split into two branches to enter the cross junction. The corresponding mean flow velocity at each of the branches (200 μm wide x 100 μm tall) immediately upstream of the cross junction was 0.17 m/s to 2.08 m/s. The microfluidic guillotine device, which consisted of 8 guillotine channels, was operated at a flow rate of 8 mL/h (referred to as regime I herein) and 36 mL/h (referred to as regime II herein), which corresponded to 1 mL/h and 4.5 mL/h per guillotine channel (i.e., 0.014 m/s and 0.063 m/s in the microfluidic channel containing the blade).^7, 9^

### Sytox Green Staining

Sytox Green staining was performed to assess the extent of wounding by the splitting process using either the hydrodynamic splitter or the microfluidic guillotine. The staining protocol was adapted from the protocol previously described.^7^ Wounded cell fragments were collected from the device outlet into a prescribed volume of fixation solution (400 μL for the hydrodynamic splitter and 250 μL for the microfluidic guillotine) into a 2 mL Eppendorf tube. The fixation solution consisted of 0.8% v/v formaldehyde (43368, Alfa Aesar) and 0.02% v/v Triton X-100 (X100-100ML, Sigma Life Sciences) in PSW at room temperature. All cell fragments were fixed at 4 seconds post-wounding by choosing appropriate lengths of outlet tubing that would transfer the cells from the outlet of the microfluidic device to the tube containing the fixation solution. The length of the outlet tubing was calculated based on the output flow rate and the diameter of the tubing. The cells were incubated in the fixing solution for 10 minutes at room temperature. The wounded cells were then stained using Sytox Green (S7020, Invitrogen) at a concentration of 2.5 μM in PSW at room temperature as follows. The fixed cells were transferred to a 4 mL glass vial with a flat bottom (C4015-21, ThermoScientific) containing 500 μL of Sytox solution using a 200 μL pipette tip with the end cut off to ensure that the diameter of the pipette tip was larger than the dimensions of the fixed cells with a cut end. The cells were incubated in Sytox solution for 30 minutes, after which 50 μL of the stained cells were washed in 500 μL of PSW in a clean 4 mL glass vial. 50 μL of the washed cells were carefully transferred onto a No. 1 glass slide for imaging using a 200 μL pipette tip with a cut end. Care was taken to avoid subsequent injury to post-wounded, fragile cell fragments. Specifically, pipette tips used for transferring cells during the fixation and staining steps were treated with 3% w/v Pluronic F-68 (J6608736, Alfa Aesar) in deionized (DI) water for at least 2 hours and then washed with PSW to prevent additional damage to the fragments. The stained cells were imaged on an EMCCD camera (Andor iXon 897, Oxford Instruments) at 15x magnification using a mercury lamp set to ND 1, and a FITC excitation/emission filter set. The mean pixel intensity of fluorescence of the stained cells was measured in ImageJ.

### Immunohistochemical staining of acetylated tubulin

To visualize the extent of cytoskeletal damage in the cells, an immunofluorescence assay was performed for acetylated tubulin localized in the km fibers in *Stentor* as previously described.^7, 33^ Briefly, cells collected at the outlet of the microfluidic devices were fixed in -20 °C methanol (250 μL of cells in 1000 μL of methanol) for 30 minutes. The cells were rehydrated at room temperature with 1:1 methanol:1xPBS mixture for 10 min, and finally in 500 μL of 1x PBS for 20 min. The cells were blocked with 2% BSA + 0.1% Triton X-100 in 1xPBS for 2 hours at room temperature or overnight at 4 °C. The cells were stained with anti-acetylated tubulin (T7451-200UL, Sigma Life Sciences) (primary antibody) diluted 1:1000 in blocking buffer for 1 hour at room temperature or overnight at 4°C, followed by 3x washes with 1x PBS. The cells were then stained with secondary antibody CFA 488 anti-mouse IgG antibody (produced in goat) (1:1000 in blocking buffer) overnight at 4 °C (SAB4600388-125UL, Fluka Analytics), followed by 3x washes in 1x PBS. 50 μL of the cell suspension was pipetted onto a No. 1 glass slide for imaging on an inverted laser scanning confocal microscope (Zeiss, LSM 780) with a 20x (NA = 0.8) objective at an excitation wavelength of 488 nm, and a broad emission filter matching the spectra of Alexa Fluor 488.

### Survival rate measurement

Survival rate was used as a measure to quantify the viability of the cells wounded by the microfluidic devices. In the current work, we quantified the *t*-day survival rate of cell fragments as the percentage of cells alive on day *t* post-cutting (*N*_*t*_) in comparison with the number of live cell fragments collected immediately after the cutting process (*N*_0_). We performed the survival assay for up to 5 days post-splitting. A cell fragment was counted to be alive if it had beating cilia or an intact plasma membrane when observed under a stereoscope. To calculate the survival rate, 200 μL of cell fragments wounded by a microfluidic device was collected in a 0.6 m long tubing. The length of the tubing was chosen such that the volume contained inside the tubing was greater than 200 μL, which ensured that all wounded cell fragments remained within the tubing after the flow inside the device was stopped. Finally, the cell fragments were ejected out of the tubing into PDMS wells (mean diameter: 12 mm, height: 10 mm) pre-filled with 400 µL of PSW. The side walls of the PDMS wells were slightly beveled to facilitate imaging of cell fragments close to the wall. The cells were ejected into the wells by inserting the free end of the tubing into the well and allowing the fluid inside the tubing to slowly drain into the well after carefully disconnecting the other end from the device outlet. To minimize further wounding of the cell fragments because of viscosity and surface tension, we ensured that the tip of the tubing was submerged in PSW inside the well. Additionally, we maintained the tubing horizontally at approximately the same level as the well so that the fragments flowed into the well gently without forming eddies. *N*_0_ was measured by counting the number of live cell fragments inside the PDMS wells immediately after cutting. To prevent the wells from drying out, they were placed inside a Petri dish with soaked paper wipes and sealed using Parafilm laboratory wrapping film (Bemis PM 996). *N*_*t*_ was calculated by counting the number of live cells in the wells on day *t* post-wounding. The regenerating fragments were not fed over the course of the survival experiments. The fragments were imaged each day until day 5 using a Canon EOS Rebel T3i DSLR camera. Images of fragments at day 0 were taken 5 – 10 minutes post-splitting.

### Visualization and image processing

To visualize the cell splitting process at the cross junction, we used a Vision Research Phantom v7.3 high speed camera mounted on a stereoscope at 5x magnification. Depending on the flow rate, the videos were recorded at frame rates ranging from 4000 to 14035 fps. The cell bisection experiments with the microfluidic guillotine were recorded on the Phantom v341 high speed camera mounted on an inverted brightfield microscope at 20 fps using a 5x objective. To characterize the performance of the hydrodynamic splitter, we evaluated cell velocity (*v*_cell_), fragments size ratio after post-splitting (*κ*), and probability of cell splitting (*P*_s_). The movies were processed using MATLAB R2019a Image Processing Toolbox. The effective cell radius, *r*_*cell*_, was estimated from a sphere with a cell volume equal to that obtained by projecting 2D image of the cell through the entire channel height, i.e., 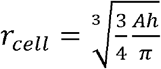, where *A* and *h*, and are the projected area of the cell and the channel height, respectively. This cell volume estimation is reasonable as the calculated *r*_*cell*_ was greater than the channel cross-sectional dimensions (100 µm × 200 µm). The velocity of the cell, *v*_cell_, was calculated by measuring cell displacement between two frames. Both *r*_*cell*_ and *v*_cell_ were calculated from images of the cells in the microfluidic device upstream of the cross junction. The splitting ratio, *κ* (≥1), was calculated by obtaining the ratio of the projected areas of the larger cell fragment to the smaller cell fragment exiting the cross junction after splitting.

### Droplet generation and splitting

We generated DI water droplets in a continuous phase of mineral oil (M5904-500ML, Sigma Aldrich, density: 840 kg/m^3^, viscosity: 0.024 Pa-s) with 2% Span 80 (85548-250ML, Fluka Analytical) as surfactant using a flow focusing microfluidic droplet generator at flow rates of 0.5 mL/h for the continuous phase and 0.5 mL/h for the dispersed phase. The flow rates for droplet generation were chosen such that the volume of each droplet generated was similar to that of a *Stentor* cell. After generation, the droplets were injected into the symmetric hydrodynamic splitter for droplet splitting using tubing connecting the outlet of the droplet generator to the inlet of the splitter. A sheath flow was used to increase the separation between consecutive droplets and modulate the droplet flow velocity during the splitting process. We performed droplet splitting at total flow rates ranging from 5.0 to 12.5 mL/h.

### Statistical analysis

Significance test was performed in MATLAB using the two-sample t-test with unequal mean and variance for the data sets. A p value smaller than 0.05 implied that the difference between the two data sets was statistically significant.

## 3. RESULTS AND DISCUSSION

### Design and operation of the hydrodynamic splitter

The hydrodynamic cell splitter consists of a microfluidic cross junction (Fig. 1A). Cells were injected into the inlet of the hydrodynamic splitter. As a cell entered the cross junction, the cell was stretched by an extensional flow until it split into two fragments (Fig. 1B). The width *w* (200 μm) of the channels that intersected to form the cross junction was chosen to be less than the diameter of a cell (∼ 200 – 400 μm) so that the cell was fully confined and centered within the channel as it entered the cross junction. In channels wider than 200 μm, cells were not always centered, which resulted in no splitting or uneven splitting. On the other hand, very narrow channels increased the likelihood of wounding the cells due to their interactions with the side walls. The fluid flow inside the cross junction was sensitive to perturbations in the pressure distribution at the outlet branches. To reduce such pressure perturbations, both outlet branches of the cross junction merged downstream to form a single outlet (See Fig. S1). Pressure shunts (Fig. 1B) were also added to the outlet branches to equalize the pressures at the two branches. The entrance fluid velocity *v*_*e*_ was controlled using a syringe pump and was varied from 0.17 m/s to 2.08 m/s, corresponding to a moderate Reynolds number *Re* of 35 to 416, respectively (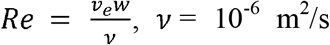 is the kinematic viscosity of water).

**Fig. 1.**
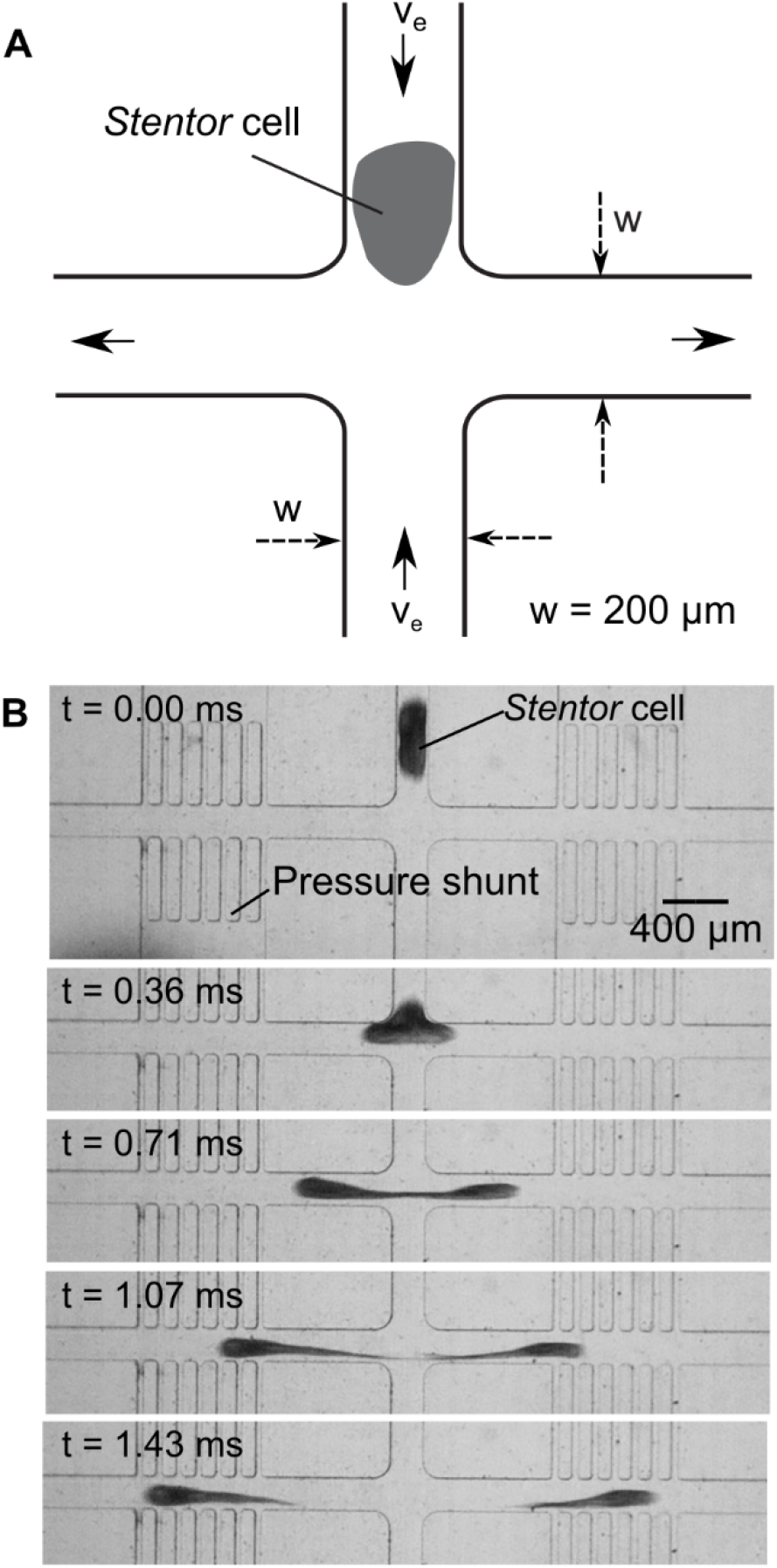
Device design and operation: **A**. Schematic diagram of the hydrodynamic splitter consisting of a cross junction. The direction of the fluid flow is indicated by the solid arrows. **B**. Time lapse showing a *Stentor* cell being split at the cross junction. The entrance velocity of the fluid is 1.39 m/s.

Fig. 1B shows a time series of a *Stentor* cell splitting at the cross junction of a hydrodynamic splitter using an entrance velocity *v*_e_ = 1.39 m/s. At the cross junction, hydrodynamic extensional stresses dominated over shear stresses and caused the cell to split into two fragments.^34^ Downstream of the cross junction, hydrodynamic shear forces could play a role in causing additional splitting of the cell fragments, but such splitting events were not observed within the field of view of our experiments. At *v*_e_ = 1.39 m/s, the splitting process took around 1 ms, estimated from the time when the cell entered the cross junction until it split into two fragments. The timescale of the splitting process varied from approximately 0.5 ms to 10 ms as the entrance velocity varied from *v*_e_ = 2.08 m/s to *v*_e_ = 0.35 m/s. Compared with the microfluidic guillotine, the timescale of splitting was 10^2^ to 10^3^ times faster in the hydrodynamic cell splitter.^9^ The high volumetric flow rate used for operating the hydrodynamic cell splitter implies that the device can process more cells in a shorter amount of time than the microfluidic guillotine. Although we only processed up to 40 cells for our device characterization, we estimate that the device could split cells at a rate of 500 *Stentor* cells per minute. This rate was about 8 times faster than the microfluidic guillotine, and 1500 times faster than manual surgery.

### Characterization of splitting probability and symmetry

Since cell splitting was driven primarily by extensional flow at the center of the cross junction,^34^ we expect the extensional stress to determine the fate of a cell (i.e., splitting or no splitting) at the cross junction. Fig. 2A shows a phase diagram of splitting events as a function of cell size and the strain rate (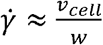, where *v*_cell_ is measured prior to the cell’s entry into the cross junction). The plot shows two distinct regimes separated by a critical strain rate of ∼3000 s^-1^, corresponding to an extensional stress, 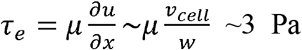 (for viscosity ∼ 10^−3^ Pa-s).^34^ Cells subjected to a strain rate higher than the critical value split at the cross junction, while those subjected to a lower strain rate deformed only without splitting.

**Fig. 2.**
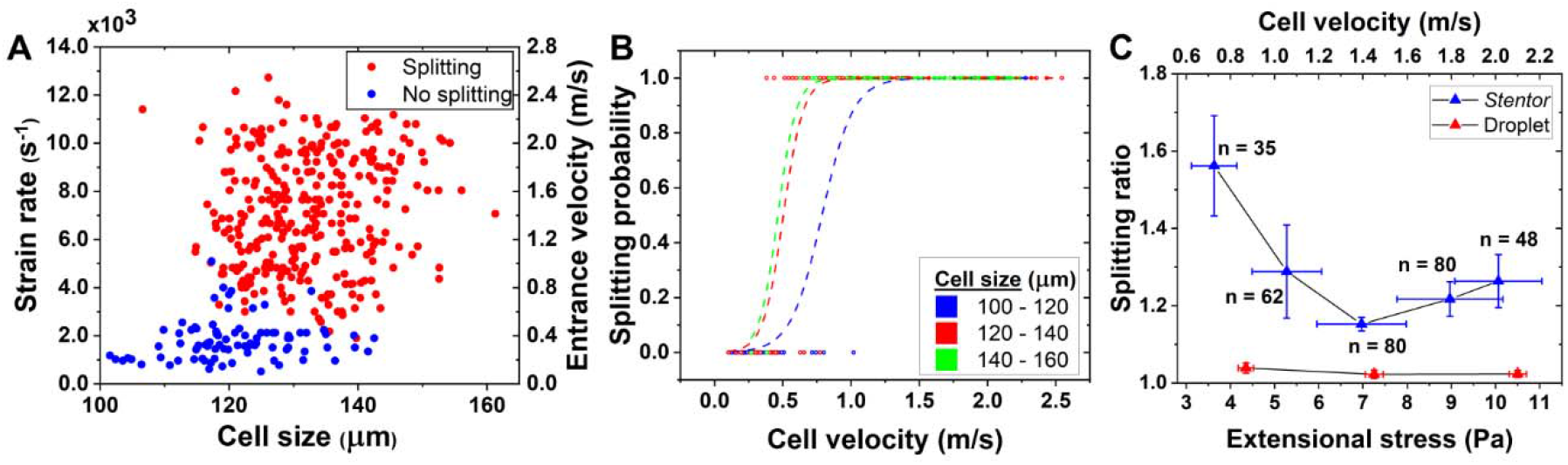
Characterization of hydrodynamic cell splitter: **A**. Phase diagram of cell splitting in the hydrodynamic splitter. The diagram indicates that there exists a critical strain rate for cell splitting. **B**. The probability of cell splitting as a function of cell velocity for different ranges of cell size. The dashed lines are the fitted logistic curves for each cell size range. **C**. Splitting ratio vs. cell velocity or extensional stress for cells or droplets. The quantity *n* represents the number of cells analyzed for each data point.

To further quantify the splitting probability, especially at strain rates close to the critical strain rate, Fig. 2B plots the splitting events and the logistic regression fits to the data (Fig.2B, dashed lines) as a function of the cell velocity for three cell size ranges (100 – 120, 120 – 140, and 140 – 160 μm). The logistic curves indicate the probability of a cell being split into two fragments. We used a sigmoid function for the splitting probability 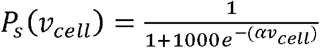 to fit the binary data. The fitting parameter α was calculated to be 8.78, 13.71, and 14.91, for cell size ranges of 100-120, 120-140, and 140-160 μm, respectively, with a coefficient of determination R^2^ values being 98.27%, 99.05% and 99.89% respectively. The individual data sets for Fig. 2B can be found in Fig. S2. The logistic curves divided the cell splitting events into three distinct regimes based on the entrance velocity. In the low entrance velocity regime (*v*_e_ < 0.5 m/s), the splitting probability *P*_s_ ≈ 0. In the high entrance velocity regime (*v*_e_ > 1.25 m/s), the splitting probability *P*_s_ ≈ 1. In these two regimes, cell splitting was independent of its size. In the intermediate velocity regime (0.5 m/s < *v*_e_ < 1.25 m/s), the splitting probability was size-dependent. The fitted curves shifted to the left with increasing cell size, indicating that, for the same entrance velocity, larger cells were more likely than smaller cells to split when the entrance velocity was between 0.5 m/s and 1.25 m/s.

Ideally, a symmetric cross junction would produce symmetric cell fragments. To quantify the symmetry of splitting, we define the splitting ratio *κ* as the ratio of the size of the larger fragment to the size of the smaller fragment. By this definition, *κ* is always greater than or equal to 1. A splitting ratio *κ* = 1 implies that the cell split symmetrically, and the two fragments were equal in size. Fig. 2C depicts the splitting ratio *κ* as a function of cell velocity *v*_cell_ and the corresponding extensional stress. We found that at low entrance velocities (i.e., low extensional stresses), the cells split asymmetrically. The degree of symmetry generally improved with increased extensional stresses. At high entrance velocities (*v*_e_ > 1.4 m/s) or extensional stresses (*τ*_*e*_≥ 7 Pa), the splitting ratio increased slightly to a value between 1.15 and 1.30. In our experiments, the cells were spaced sufficiently far apart (usually a cell length or more) so that the split cells did not interfere with subsequent splitting. To assess whether the asymmetry of the cell splitting arose from the flow field or from the inherent inhomogeneity of the cellular structures, we quantified the splitting ratio for the splitting of DI water droplets in a continuous phase of mineral oil with 2% Span 80 as surfactant. The flow velocity was chosen such that the extensional stresses for droplet splitting were similar in magnitude to those experienced by the cells. We found that droplet splitting was more symmetric than cell splitting, with splitting ratios very close to 1 (*κ =* 1.02 – 1.04).

There are several possible reasons for asymmetric cell splitting inside a symmetric cross junction. First, the morphology and the internal structure of the cell is inherently inhomogeneous.^15^ The presence of km fibers, macronuclear nodes and other organelles in *Stentor* likely led to local variations in the stiffness and deformability of the cell. Therefore, even though the flow was symmetrical, the cell did not split equally. At high flow rates when the hydrodynamic stresses were sufficiently large, the effect of the cellular inhomogeneities may have been reduced, possibly due to shear thinning effects of the cytoplasm and/or the organelles,^35-37^ resulting in more symmetric splitting and lowering of the splitting ratio. The rheological properties of cellular cytoplasm are known to be shear rate dependent, and the cytoplasm has been found to be shear thinning in some cell types, including human neutrophils^35^ and in the single-cell protozoan *Entamoeba histolytica*.^37^ Additionally, suspensions of F-actin and microtubules, important constituents of the cell cytoskeleton, are known to be shear thinning.^36^ The rheology of cytoskeletal components, mainly the microtubule ribbons in the km fibers in *Stentor*, could give its cytoplasm shear thinning characteristics.

Second, streamwise vortices are known to develop in water downstream of the cross junction at moderately high Reynolds numbers (onset *Re* = 107 for a channel height to width ratio of 0.45) comparable to those used in the current study.^38^ The presence of such asymmetric vortices could lead to asymmetric splitting of cells. When cell splitting was performed at a Reynolds number of 346 (*v*_e_ = 2.08 m/s), we noticed that in some cases, there was flapping of the “tail” region of the fragment immediately following the split (Movie S1). This flapping of the tail of the fragment could be indicative of the presence of asymmetric vortices in the flow. We did not observe such flapping or vortices in droplet splitting experiments, because they were performed at low Reynolds numbers (Re ∼ 0.1). The reason why the splitting ratio increased again at *v*_e_ > 1.4 m/s is under current investigation. Nevertheless, despite the above factors, the overall splitting ratio was still relatively small (*κ* <1.56), and we believe the symmetry was sufficient for most downstream biological applications.

### Characterization of cellular wounding and viability post-splitting

Although our device was capable of splitting cells at a very high speed, the high hydrodynamic stresses imposed on the cells could affect the extent and nature of cellular wounding and subsequent viability. Fig. 3A compares the wounds created by the hydrodynamic splitter against those created by the microfluidic guillotine. Sytox Green staining was performed to assess the extent of wounding caused by each of these devices. Sytox Green is a nucleic acid stain which does not penetrate intact plasma membranes, but enters cells with compromised membranes, where it generates a fluorescence signal upon binding with nucleic acids. The signal intensity tends to increase with the severity of the wound and therefore could be a qualitative indication of the degree of wounding.^7, 9^ For all wounded conditions shown, the measurements corresponded to cells fixed at 4 seconds post-splitting.

**Fig. 3.**
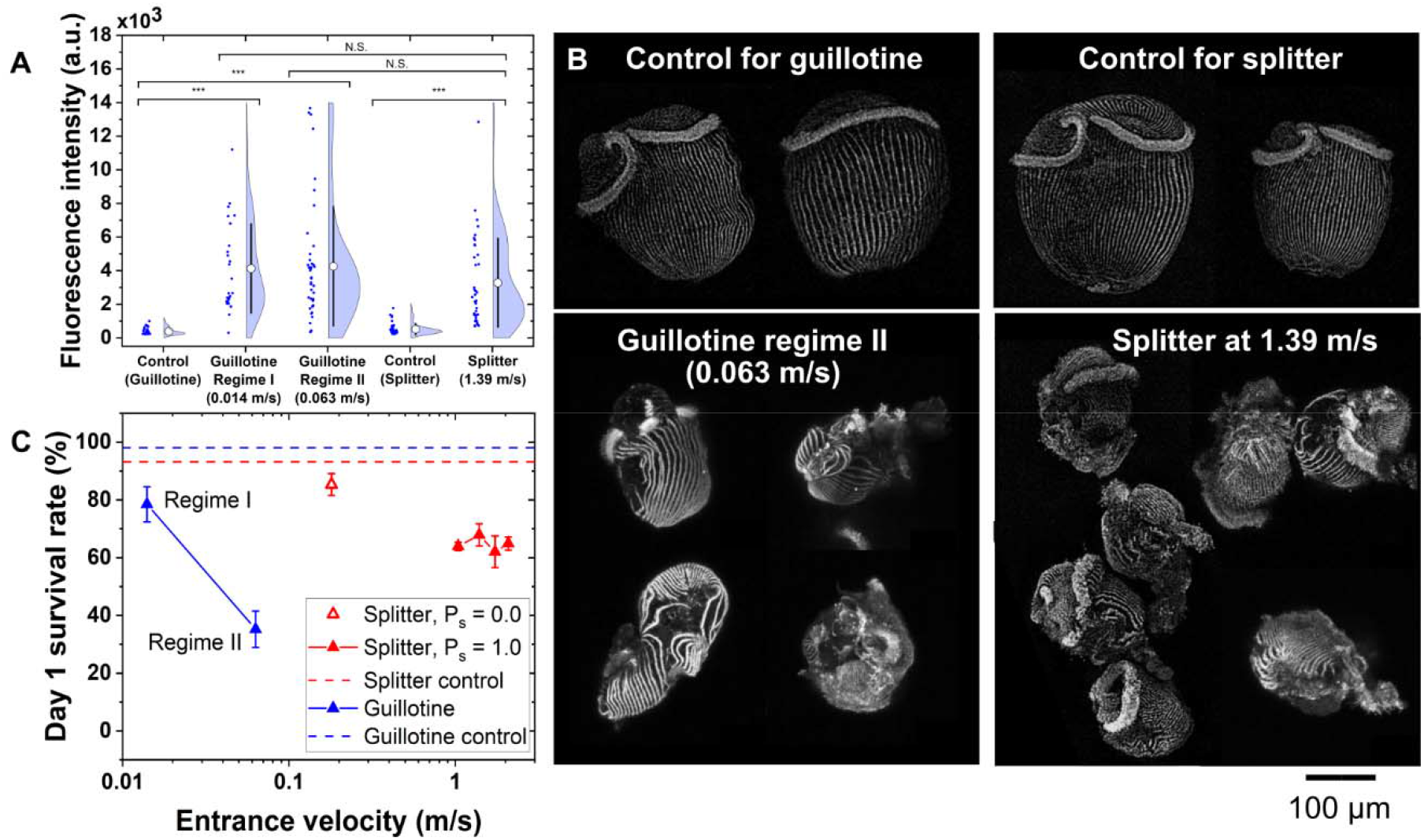
Characterization of cellular wounding and viability post-splitting. **A**. Fluorescence intensity of Sytox Green stained cells that were fixed 4 seconds post-wounding. (\*\*\**p* < 0.001, N.S: not significant). **B**. Fluorescence images of *Stentor* cells labeled with anti-acetylated tubulin show the orientation of km fibers under different conditions. The km fibers are intact in control cells, but are discontinuous in wounded cells. The images have been adjusted for brightness and contrast for the purpose of visualization. **C**. Day 1 survival rate vs. flow velocity for the guillotine and the hydrodynamic splitter (for splitting probability *P*_s_ = 0.0 and 1.0, respectively). The survival rate for control cells for the two devices are shown in dotted lines. The variance in the survival rate is shown in terms of standard error.

The control for each device consisted of flowing unwounded *Stentor* cells through a piece of tubing (inner diameter of 762 µm) directly into the fixing solution at the same flow rates as those used for collecting cell fragments from the microfluidic guillotine or the hydrodynamic splitter after splitting. This control was intended to examine whether the outlet flow would further wound the cell fragments. For the guillotine control, the *Stentor* cells flowed in the tubing at a rate of 18 mL/h which corresponded to a mean flow velocity of 0.01 m/s in the tubing. This flow rate matched the outlet flow rate used for collecting cell fragments from the tubing after splitting the cells with the microfluidic guillotine operated in regime I. At this flow velocity, the unwounded *Stentor* cells were found to exhibit a low fluorescence signal that was significantly lower than the fluorescence signal exhibited by cells wounded by the guillotine operated in regime I (*p* = 2.68×10^−7^) and regime II (*p* = 8.98×10^−10^). For the hydrodynamic cell splitter control, the cells flowed at a rate of 200 mL/h, which corresponded to a mean flow velocity of 0.12 m/s in the tubing. At this flow velocity, the unwounded *Stentor* cells had slightly higher fluorescence signals than the guillotine control. The difference between the two control conditions was statistically significant with *p* = 0.0375. However, the Sytox Green intensity for both control conditions remained significantly lower than cells wounded by the splitter (*p* = 9.25×10^−10^). The slight difference in the two control sets could be because of the higher shear stress that unwounded cells might experience inside the tubing at the higher flow rate. The higher shear stress could create pores on the cell membrane which allowed Sytox Green to enter the cell. However, these pores were not sufficient to create drastic wounds, evident from the large difference in signal intensity between the control set and the split cells.

The average fluorescence intensity corresponding to the two flow conditions of the microfluidic guillotine were 3900 a.u. (arbitrary intensity units) in regime I, and 4300 a.u. in regime II. In comparison with the guillotine, cells split using the hydrodynamic splitter at *v*_e_ = 1.39 m/s were found to exhibit a lower average fluorescence intensity of 3000 a.u.. However, the large variance in the data for the two devices meant that cells wounded by the hydrodynamic splitter were not significantly different from cells wounded by the guillotine (*p* = 0.080 at regime I and *p* = 0.052 at regime II). The data sets for each condition of Sytox Green staining can be found in Fig. S3.

Fig. 3B shows images of tubulin-stained cells marking the tubulin-rich km fibers. Wounded *Stentor* cells were fixed within 10 seconds post-wounding. The unwounded control cells showed intact ribbons of km fibers running along the length of the cell body, indicating that the cells were unlikely to be wounded. In comparison, cells fixed post-bisection in the guillotine and the hydrodynamic splitter exhibited significant wounding. The km fibers appeared discontinuous at multiple locations. Qualitatively, there appeared to be no significant difference in the extent of wounding between the two devices, which is consistent with the Sytox Green data.

Fig. 3C compares the 1-day survival rate of cells bisected by the microfluidic guillotine versus the hydrodynamic splitter. The dotted lines indicate the 1-day survival rates of the corresponding controls. The hydrodynamic splitter recorded a survival rate of 86% for *v*_e_ = 0.17 m/s. At this flow rate, the cells deformed but did not split (i.e., splitting probability *P*_s_ = 0.0). Also, the compressive stress on the cell 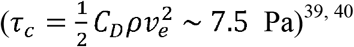 was larger than the extensional stress (*τ*_*e*_ ∼ 0.85 Pa). The fact that the survival at this flow rate was lower than the control indicates that compression at the cross junction was sufficient to create wounds which resulted in the death of some cells, even though the extensional stress did not lead to splitting at this velocity. In the high flow velocity regime (*v*_e_ ≥ 1.04 m/s) where all cells split (i.e., *P*_s_ = 1.0), the survival rate dropped to ∼60% and remained between 60% and 65% from *v*_e_ = 1.04 m/s to *v*_e_ = 2.08 m/s. At these velocities, the splitting was relatively symmetric (*κ* ∼ 1.15 - 1.30). Given that the hydrodynamic shear and extensional stresses increased by a factor of 2 from *v*_e_ = 1.04 m/s to 2.08 m/s, these results indicate that increased hydrodynamic stresses did not significantly affect 1-day cell viability at these velocities and corresponding stresses.

In comparison, the survival rate for cells cut using the guillotine was ∼78% in regime I and 35% in regime II. The guillotine in regime I had higher cell viability than the hydrodynamic splitter, but the guillotine in regime II had lower viability than the hydrodynamic splitter, even though they had similar extents of cellular wounding as indicated by the Sytox Green assay and the anti-acetylated tubulin staining. The reason for the low viability of cells cut in regime II was likely due to the large number of small cell fragments generated during the cutting process (described in more detail below).

### Asymmetric splitting of cells

The hydrodynamic cell splitter was designed to ensure that the flow rates in the two outlet branches were equal. This symmetrical flow distribution was achieved by means of pressure shunts. The symmetrical flow distribution enabled symmetrical splitting of homogenous droplets, and slightly asymmetrical splitting of constitutively inhomogeneous cells (*κ* = 1.15 – 1.56). By making small adjustments to the geometry of the device, it is possible to control the ratio of the flow rates in the left and right outlet branches of the cross junction, which would enable us to modulate the splitting ratio of *Stentor* cells at the cross junction, and thereby generate smaller cell fragments.^41^ Fig. 4A shows a schematic diagram of the asymmetric hydrodynamic cell splitter designed to split cells asymmetrically at the cross junction. To induce asymmetric splitting of the cells, the hydrodynamic splitter was modified by removing the pressure shunts and introducing a subsidiary flow channel on the right outlet branch. A subsidiary flow entered the right outlet branch at a velocity *v*_s_. This additional flow introduced an asymmetry in the flow field at the cross junction. Numerical simulation showed that the subsidiary flow caused the main flow to divide unevenly at the cross junction, with a larger proportion of the main flow exiting the cross junction to the left. An increase in subsidiary flow velocity increased the proportion of the main flow exiting the cross junction to the left compared to the right (Fig. S4D). The uneven distribution of flow rates is known to create asymmetric splitting of droplets,^41^ and is also likely to induce asymmetric splitting of the cells. Additionally, the asymmetric splitter contained a constriction on one side of the cross junction. Numerical simulation of the flow at a cross junction showed that the presence of a constriction increased the extension rate in the fluid flow (Fig. S4C). Experimentally, we observed that the addition of the constriction increased the localized stretching of the cells and increased the probability of asymmetrical splitting of the cells, especially at high values of *v*_s_ (Fig. S5A). We found that the splitting ratio was relatively insensitive to the constriction width (Fig. S5B).

**Fig. 4.**
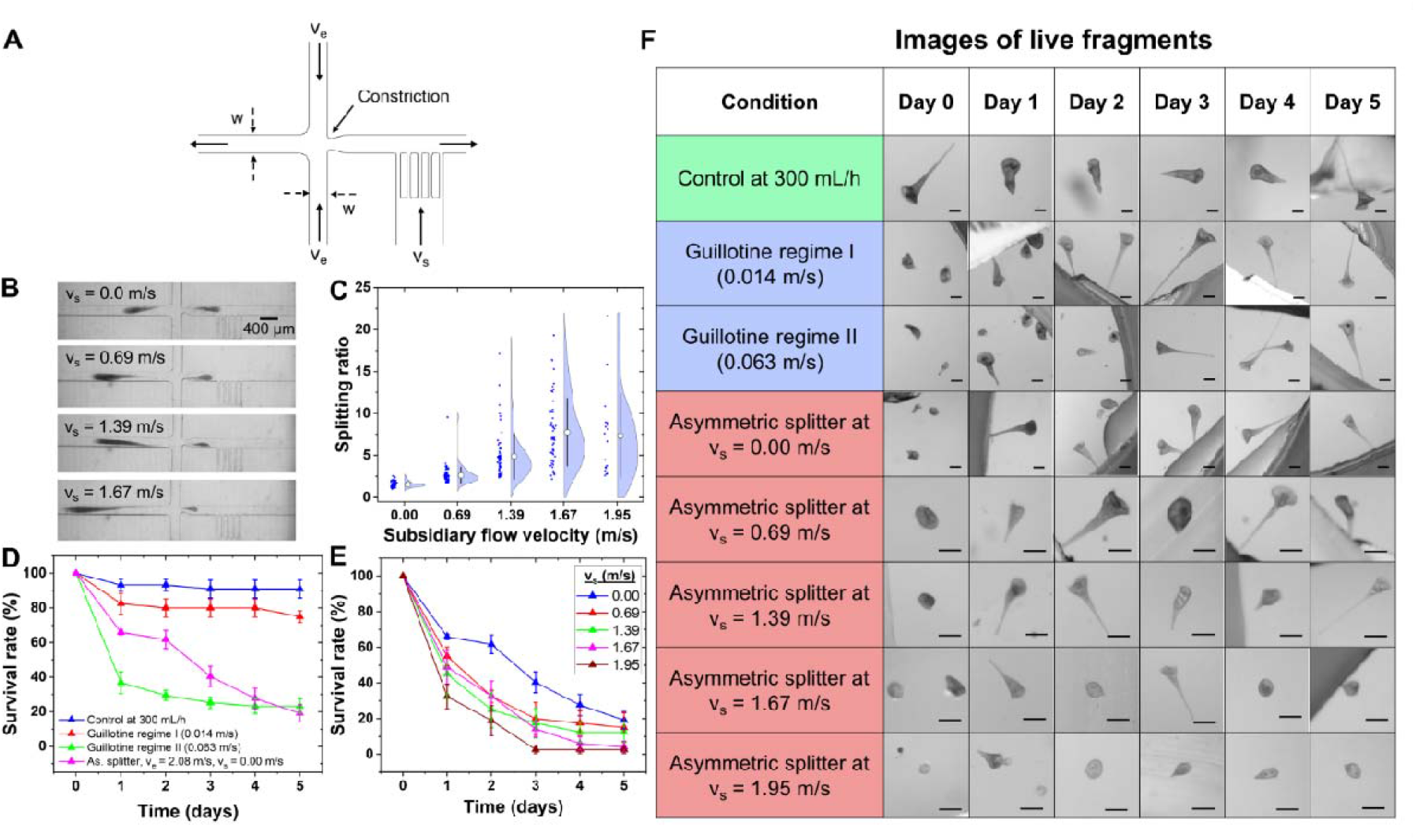
Characterization of the asymmetric hydrodynamic splitter. **A**. Schematic diagram of the device. The asymmetric splitter has an 80 µm constriction on the right outlet branch of the cross junction. **B**. Representative images showing cell splitting at the cross junction for different subsidiary flow velocities. The entrance velocity was maintained constant at 2.08 m/s in all cases. **C**. Splitting ratio vs. subsidiary flow velocity *v*_s_. **D**. 5-day survival rate for the guillotine and the asymmetric splitter without subsidiary flow velocity. **E**. 5-day survival rate for the asymmetric splitter for different subsidiary flow velocities. In both D and E, the variance is shown in terms of the standard error. **F**. Table showing representative images of healthiest-looking cell fragments over 5 days for different conditions. Scale bars in all images represent 125 µm. See Fig. S7 for table showing the unhealthy-looking cell fragments.

Fig. 4B shows representative images of asymmetric cell splitting for different subsidiary flow velocities. The main flow velocity was held constant at *v*_e_ = 2.08 m/s. The subsidiary flow velocity *v*_s_ varied from 0 to 1.67 m/s. The larger fragment of the split cell always flowed into the left outlet branch while the smaller fragment always flowed into the right branch. The two fragments were collected separately at two separate outlets.

Fig. 4C shows that the splitting ratio *κ*, i.e., the size of the larger fragment in the left outlet branch relative to the size of the smaller fragment in the right branch, generally increased with increasing *v*_s_ from 0 to 1.67 m/s. Further increase in subsidiary flow velocity beyond *v*_s_ = 1.67 m/s did not increase the average splitting ratio. When no subsidiary flow was applied, the splitting ratio was 1.31, which is similar to the splitting ratio for the symmetric splitter (Fig. 2C). At *v*_s_ = 1.67 m/s, the average splitting ratio was estimated to be *κ* = 7.5, which indicates that the smaller fragment was approximately 1/8.5^th^ the size of the original cell, based on projected area. In some cells, our device achieved splitting ratios higher than 7.5, although the frequency of occurrence of these ratios was low. We also observed an increase in the variance of the splitting ratio as *v*_s_ increased, likely due to the increased sensitivity to the local distribution of cellular components at the site of splitting.

We note that one of the limitations of the parameter *κ* as defined in this work is that it is estimated from the projected area of the fragments exiting the cross junction, assuming the cell fragment occupied the entire height of the channel. As the splitting ratio increased, it was likely that the small fragments did not occupy the entire height of the channel. Therefore, our method to measure the size of cell fragments likely overestimated the actual size of the fragments.

Figs. 4D – E describe the variation in the survival rate of the cell fragments over a period of 5 days post-splitting. For the asymmetric splitter, we only collected the fragments exiting the cross junction through the right outlet containing the constriction, i.e., the smaller cell fragment. To draw a baseline for comparison of the data, we measured the survival rate of a control set. The control set consisted of cells flowing through a tubing (inner diameter 762 µm) at 300 mL/h (corresponding to a mean flow velocity 0.18 m/s in the tubing), which was the flow rate used for generating the main flow in the asymmetric splitter. This flow condition was chosen as the control to emulate the hydrodynamic stresses that the unwounded cells experience due to the flow inside the tubing prior to entering the microfluidic device.

The survival rate for the control set was higher than 90% and was found to stay nearly constant over 5 days. The top row in Fig. 4F shows representative images of cells taken from the control set for 5 days. The cells were large (body length > 250 µm), and appeared to be generally healthy, indicated by the trumpet shape, the presence of an oral apparatus, intact plasma membrane and beating cilia. These cells could be observed either in an extended shape adhering to the solid surface at the holdfast, or swimming freely in the medium.

Fragments generated from the guillotine in regime I were found to regenerate into functioning cells (Fig 4F, 2^nd^ row), which appeared to closely resemble cells from the control set. The mean survival rate of the fragments was measured to be 82% on day 1 (Fig. 4D). The survival rate stayed nearly constant afterwards. These data are consistent with the relatively gentle splitting of the cells in regime I, and most of the cell fragments were able to regenerate to healthy-looking cells.

The guillotine operated in regime II produced a large number of small fragments, about three times the number of cells that were injected into the device. These small fragments were generated either as the cell was cut at the guillotine, or from further splitting of the cell fragments downstream of the guillotine. There was a large variation in the size and morphology of fragments produced (Fig 4F, 3^rd^ row). Many of the fragments appeared as transparent spheres with membrane-bound beating cilia (Movie. S2). The small transparent spherical fragments were usually short-lived, with most of them not remaining viable past day 1. These short-lived fragments contributed to the large decrease (64%) in the mean survival rate between days 0 and 1 for fragments produced from the guillotine in regime II (Fig. 4D). The decrease in survival rate was less steep over the next 4 days and saturated at about 23%. The cell fragments that stayed viable on day 5 appeared to be healthy and resembled cells from the control set.

In comparison with the guillotine operated in regime II, the asymmetric splitter operated at *v*_e_ = 2.08 m/s and *v*_s_ = 0 m/s (i.e., symmetric splitting) produced approximately 25% more cell fragments than the number of cells that were injected. The proportion of short-lived transparent fragments in the medium after splitting was smaller than those from the guillotine operated in regime II. Therefore, the asymmetric splitter without any subsidiary flow recorded a 34% decrease in survival rate between days 0 and 1 post-splitting, which was about half of the drop recorded by the guillotine in regime II (Fig. 4D). However, the survival rate dropped continuously to 19% at the end of day 5, which is slightly less than the survival rate of the cell fragments generated by the guillotine in regime II. At the end of day 5, the cells that survived appeared to be healthy and their morphologies were similar to that of the control cells.

In the case of the microfluidic guillotine, where the cell fragments experienced lower hydrodynamic stresses, the decrease in survival rate between days 1 and 5 was 7% (for regime I, operated at 0.014 m/s) and 13% (for regime II, operated at 0.063 m/s). Over the same time period, the decrease in survival rate was 47% when the hydrodynamic splitter was operated at v_e_ = 2.08 m/s and *v*_s_ = 0.0 m/s (i.e., symmetrical splitting). This result suggests that even though the extent of wounding immediately (4 seconds) post-splitting appeared similar in both the guillotine and hydrodynamic splitter as measured by the Sytox Green assay and immunohistochemical staining of the km fibers (Fig. 3A), increased hydrodynamic stress experienced by *Stentor* cells likely affects their long-term repair and regeneration, a phenomenon which has not been previously reported.

To further characterize the effect of hydrodynamic splitting on cell survival, Fig. 4E shows the survival rates of fragments collected from the asymmetric splitter operated under different subsidiary velocities. We maintained the entrance velocity constant at *v*_e_ = 2.08 m/s and collected only the smaller portion of the split cells that exited the right branch of the cross junction. The survival rates were found to decrease over the course of 5 days post-splitting for all flow conditions. The survival rate also tends to decrease with increasing subsidiary flow velocity, i.e., decreasing cell fragment sizes. Cells split at *v*_s_ = 0 m/s had the highest survival rate, with a 66% survival rate on day 1 and 19% on day 5. On the other hand, cells split at *v*_s_ = 1.95 m/s had the lowest survival rates, with 32% on day 1 and 3% on day 5. Although the survival rate of cells split by the asymmetric splitter at *v*_s_=1.67 m/s appeared higher than those at *v*_s_=1.39 m/s on days 1 and 2, they were not statistically different due to the large variance in splitting ratio observed for cells split at higher subsidiary flow velocities (Fig. 4E). Qualitatively, *Stentor* cells split by the asymmetric splitter operated at high subsidiary flow velocities displayed a range of morphologies (Fig. 4F, bottom 3 rows). A few cell fragments appeared to have regenerated the normal trumpet shape, though they were smaller than those in the unwounded control set. A fraction of the cells had a spherical morphology, as discussed earlier. Many of these fragments appeared to be transparent and deficient of pigmentation.

The decrease in survival with increasing subsidiary flow velocity may be attributed to both the decrease in the size of the cell fragments, and the increased hydrodynamic stress the smaller cell fragments were exposed to. Regarding the reduced viability in smaller cell fragments, our results are consistent with previous work performed by Tartar and Lillie, who studied the effect of fragment size on regeneration.^15, 16^ The presence of nuclear nodes is a necessary but insufficient condition for full regeneration.^16^ Nucleated fragments (i.e., fragments containing at least one macronuclear node) below a critical size (∼ 80 µm) were reported to be incapable of full regeneration.^15, 16^ Additionally, nucleated fragments above the critical size require sufficient cellular materials for regeneration. Previously, Tartar has shown that *Stentor coeruleus* cells with a single macronuclear node having a cytoplasmic volume of 1 to 6 times that of a normal cell show either delayed regeneration of the oral apparatus or no regeneration at all.^20^ In that study, hypo-nucleate *Stentors* were created by excising all but one macronuclear node from a healthy cell by manual surgery and grafting the cell with one or more cells where all macronuclear nodes had been removed. This process created “chimera” cells which had a nucleocytoplasmic ratio ranging from 400 to 6000, in comparison with 126 to 387 in a normal healthy *Stentor*.^16, 20^ Lillie investigated regeneration in *Stentor polymorphus* cell fragments with nucleocytoplasmic ratios lower than those in normal healthy cells. He found that naked macronuclear nodes (20 – 25 µm) without cytoplasm were incapable of regeneration, and cell fragments (with a size up to 69 µm) containing multiple macronuclear nodes (up to 7) but insufficient cytoplasm were unable to regenerate.^16^ The lack of regeneration in very low nucleocytoplasmic ratio conditions is likely due to insufficient cellular components (e.g., mitochondria) essential for regeneration. In our experiments, the smaller cell fragments were more likely to have an unfavorable nucleocytoplasmic ratio, therefore leading to reduced long term viability.

The reduced viability in small fragments could also be due to their exposure to increased hydrodynamic stress. Many biological systems, including human cells, are responsive to hydrodynamic shear. For example, vascular and pulmonary endothelial cells respond to hemodynamic shear stress through stretch-mediated activation of mechanically gated ion-channels.^42-45^ Single-cell organisms such as dinoflagellates exhibit bioluminescence in response to hydrodynamic shear stress.^46^ In dinoflagellate *Lingulodinium polyedrum*, hydrodynamic stresses are believed to increase the fluidity of the plasma membrane thereby activating GTP binding proteins, as well as to cause an influx of Ca^2+^ ions into the cytoplasm leading to a flow of protons into the cytoplasm from vacuoles to trigger bioluminescence.^46^ Hydrodynamic stresses are also known to damage cells and reduce their viability, but the extent of damage depends on the magnitude of shear stresses and the cell type.^47-50^ For adherent mammalian cells, mechanical forces are transmitted from the integrins to the nucleus via the cytoskeletal fibers to trigger signaling pathways and can lead to cell death.^51^ Apoptosis and autophagy were induced in cancer cells when subjected to laminar shear stress (0.05 – 1.2 Pa) for an extended period of time (12 – 72 hours) through the BMPRIB/Smad1/5/p38 MAPK signaling pathway.^52^ Less is known about the effect of hydrodynamic shear on ciliates. However, a previous study showed that the use of Pluronic F-68 as a cell membrane stabilizer against hydrodynamic shearing succeeded in increasing cell viability in the single-cell ciliated protozoan *Tetrahymena thermophila*.^53^

In our asymmetric splitter, the smaller cell fragments were exposed to a strain rate of 10 × 10^3^ s^-1^ to 20 × 10^3^ s^-1^, corresponding to stresses of 10 – 20 Pa. Although the cells were exposed to high stress for only ∼40 – 500 ms when they were inside the microchannel, it is likely that the large magnitude of hydrodynamic stresses adversely affected the *Stentor* cells both for unwounded cells and cell fragments, which could lead to altered cellular behavior and reduced long-term viability. We observed multiple instances (7 cannibals in 31 sets of splitting experiments at *v*_e_ = 2.08 m/s, with 3 – 24 cell fragments in each set, i.e., a total of 93 – 744 cell fragments) of *Stentor* cells exhibiting cannibalism in the survival assays where cells were split at an entrance velocity of *v*_e_ = 2.08 m/s, and one instance in the control set for the hydrodynamic splitter where the cells were subjected to a velocity of 0.18 m/s inside the tubing (Fig. S6). Cannibalism was not observed in survival assays for the guillotines or for the hydrodynamic splitter at entrance velocities *v*_e_ < 2.08 m/s. Cannibalism has been reported in other cell types, including human cancer cells.^54, 55^ Although cannibalism in single-cell protozoa,^15, 56-58^ including *Stentor*, has been reported in the literature, the conditions leading to cannibalism in *Stentor* are not well understood.^15, 58, 59^ It is known, however, that starvation does not drive cannibalism in *Stentor*.^15^ Indeed, even though our cells were not fed for five days during all 5-day survival assays, we did not observe any *Stentor* cannibalism in the guillotine experiments which were performed at low flow velocities. An increased occurrence of cannibalism at high entrance velocities appears to suggest that hydrodynamic stresses could be a driving factor of this altered cell behavior. We note that for the survival experiments, we excluded experiments where cannibalism was observed, so cannibalism itself does not explain the decrease in survival rates in the hydrodynamic splitter. Increased metabolic stress due to hydrodynamic stresses could explain the decrease in survival rate of cell fragments generated by the hydrodynamic splitter over the course of 5 days. However, the effect of hydrodynamic stress and the corresponding signaling pathways have not been reported in *Stentor*. Our observations indicate a previously undescribed mechanosensitive pathway whereby hydrodynamic stress impacts the long-term repair in *Stentor*. Further investigation of this pathway is the subject of a separate study.

## 4. CONCLUSIONS

In this paper, we described the hydrodynamic splitter as a microfluidic tool for splitting cells in a non-contact manner using extensional flow at a cross junction. This device offers a new way to split *Stentor* cells at a high throughput of ∼500 cells/minute without the accumulation of cellular debris. Cell splitting is independent of cell size when the device operates at a sufficiently high flow rate. The fragments generated by the symmetric hydrodynamic splitter showed a small degree of asymmetric splitting *κ* ∼ 1.15 – 1.56, which is likely caused by the inherent inhomogeneities within the cell. Cell survival assays indicate that the fragments generated after splitting had a 1-day survival of ∼60%. The extent of wounding in the hydrodynamic splitter is similar to that created by the microfluidic guillotine operated in regime II. By introducing a subsidiary flow that results in asymmetric splitting, it is possible to generate fragments as small as ∼8.5 times smaller than the original cell size. A 5-day survival assay showed that the viability of cell fragments decreased with increasing subsidiary flow velocity and also with time. The trend likely arises from the fact that increased subsidiary flow decreases cell fragment size and increases the hydrodynamic stress on the fragments. Small fragments may lack sufficient nuclear and/or cytoplasmic materials for full regeneration and survival. Increased hydrodynamic stress, albeit for a very short period of time (<1 second), may induce undesirable cellular stress altering cell behavior and survival. Nevertheless, we believe that our device can be useful for generating fragments at a high throughput manner for studying phenomena where long-term viability is not critical, such as the closure of plasma membrane wounds which occurs on the time scale of ∼100-1000 seconds in *Stentor*, or for dissecting cellular systems that maybe more robust against mechanical stress if long-term viability is important. Finally, our study inadvertently uncovered a previously unknown effect of hydrodynamic stress on the long-term repair process in *Stentor*. Further investigation is needed to elucidate the biological mechanisms of how mechanical stress, both the magnitude and duration of exposure, determines different stages of cellular repair and viability. As the hydrodynamic splitter applies high hydrodynamic stresses on soft samples until mechanical failure, the device could be applied for dissociating small tissues or organoids into cell clusters or single cells, and also for measuring the limits of mechanical failure in soft materials, including biological samples.

## Conflicts of Interest

There are no conflicts to declare.

## Acknowledgments

The authors would like to thank Dr. Ambika V. Nadkarni (Department of Mechanical Engineering, Stanford University) and Dr. Wallace F. Marshall (Department of Biochemistry and Biophysics, University of California San Francisco) for their valuable feedback on the work. The work was supported by the National Science Foundation (NSF Award: 1938109), and in part by the Center for Cellular Construction, which is a Science and Technology Center funded by the National Science Foundation (NSF Award: DBI-1548297). Device fabrication was performed at the Stanford Nano Shared Facilities (SNSF), supported by the National Science Foundation under award ECCS-2026822.

## Notes

### Competing Interest Statement

The authors have declared no competing interest.

