## Supplementary Information for "Hydrodynamic dissection of single cells in a microfluidic system"

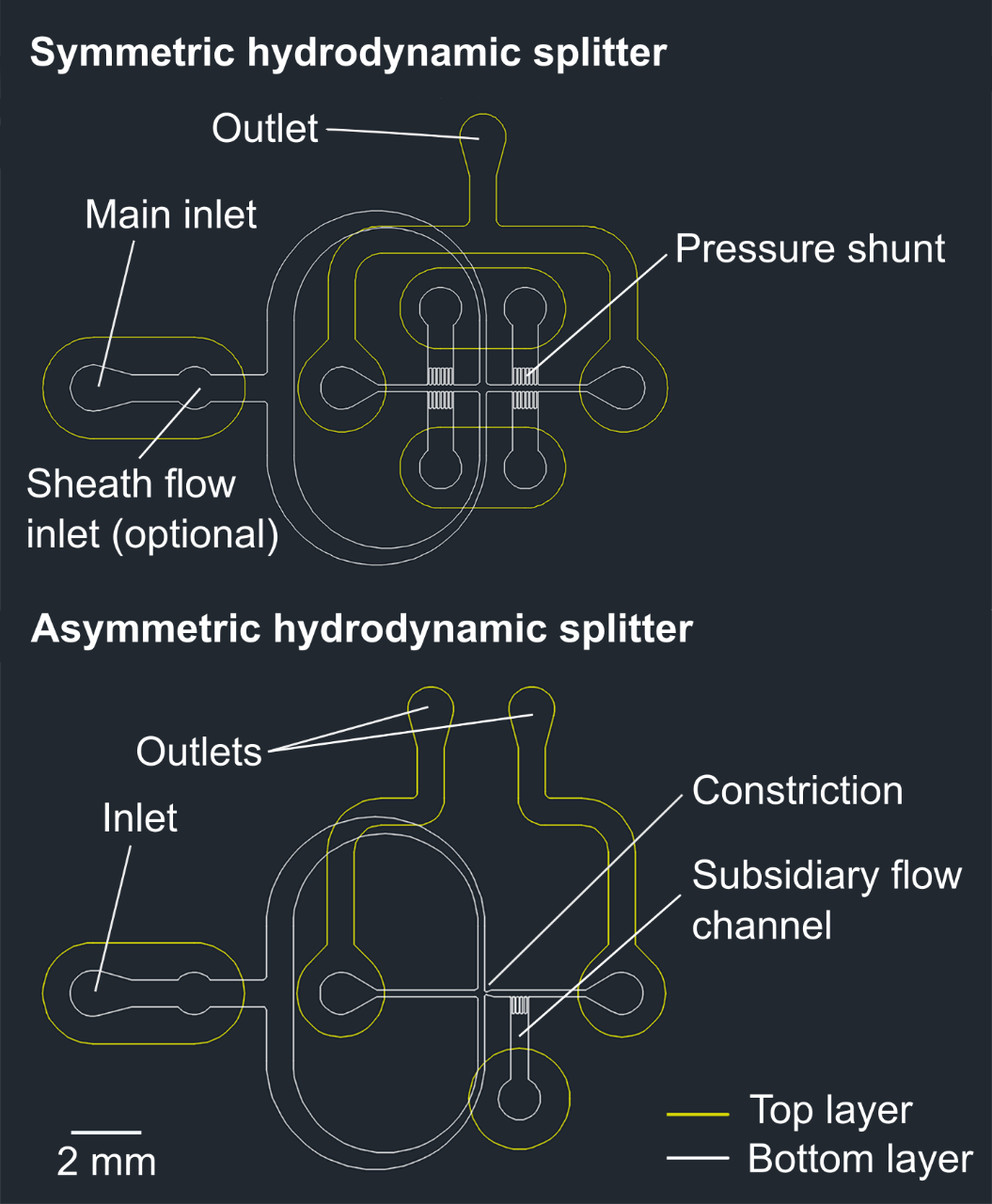

**Fig. S1. Schematic diagrams of the hydrodynamic splitters.** The device consists of two layers of channels. The features on the top and bottom PDMS layers are indicated by yellow and white lines, respectively. The symmetric splitter consists of an inlet and an outlet. Pressure shunts at the two exiting branches equalize pressure to ensure symmetric splitting. The asymmetric splitter consists of an inlet and two outlets. A subsidiary flow channel and a constriction are included to enable asymmetric splitting. In both devices, there is an inlet for sheath flow. This inlet was only used for droplet splitting with the symmetric splitter.

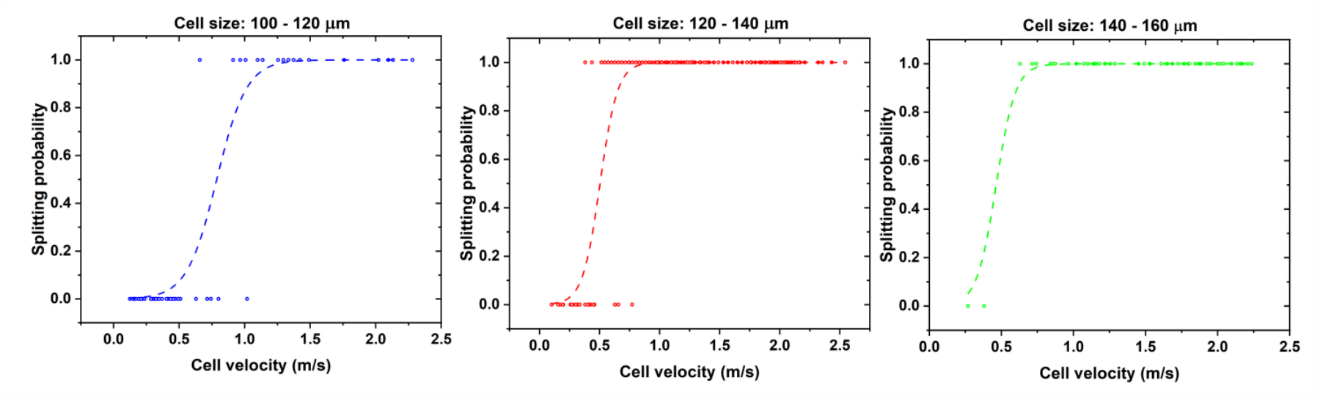

**Fig. S2.** Data sets for the splitting probability as a function of cell velocity is shown for three ranges of cell sizes: 100 – 120 µm, 120 – 140 µm and 140 – 160 µm. The dashed lines represent the fitting logistic curve for each data set.

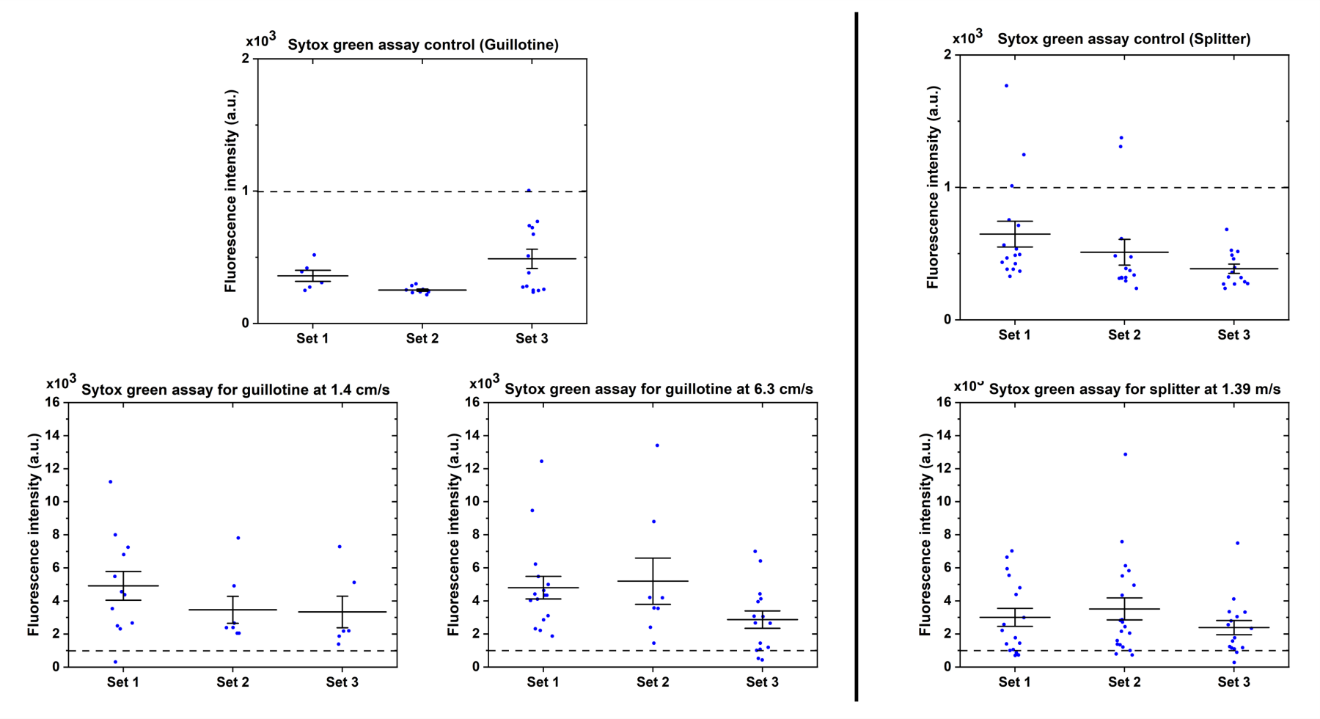

**Fig. S3. Data sets for Sytox Green staining experiments.** The data sets for the microfluidic guillotine are on the left, and the hydrodynamic splitter are on the right. The threshold fluorescence for unwounded control experiments was set based on the guillotine control sets and is denoted by a dotted line.

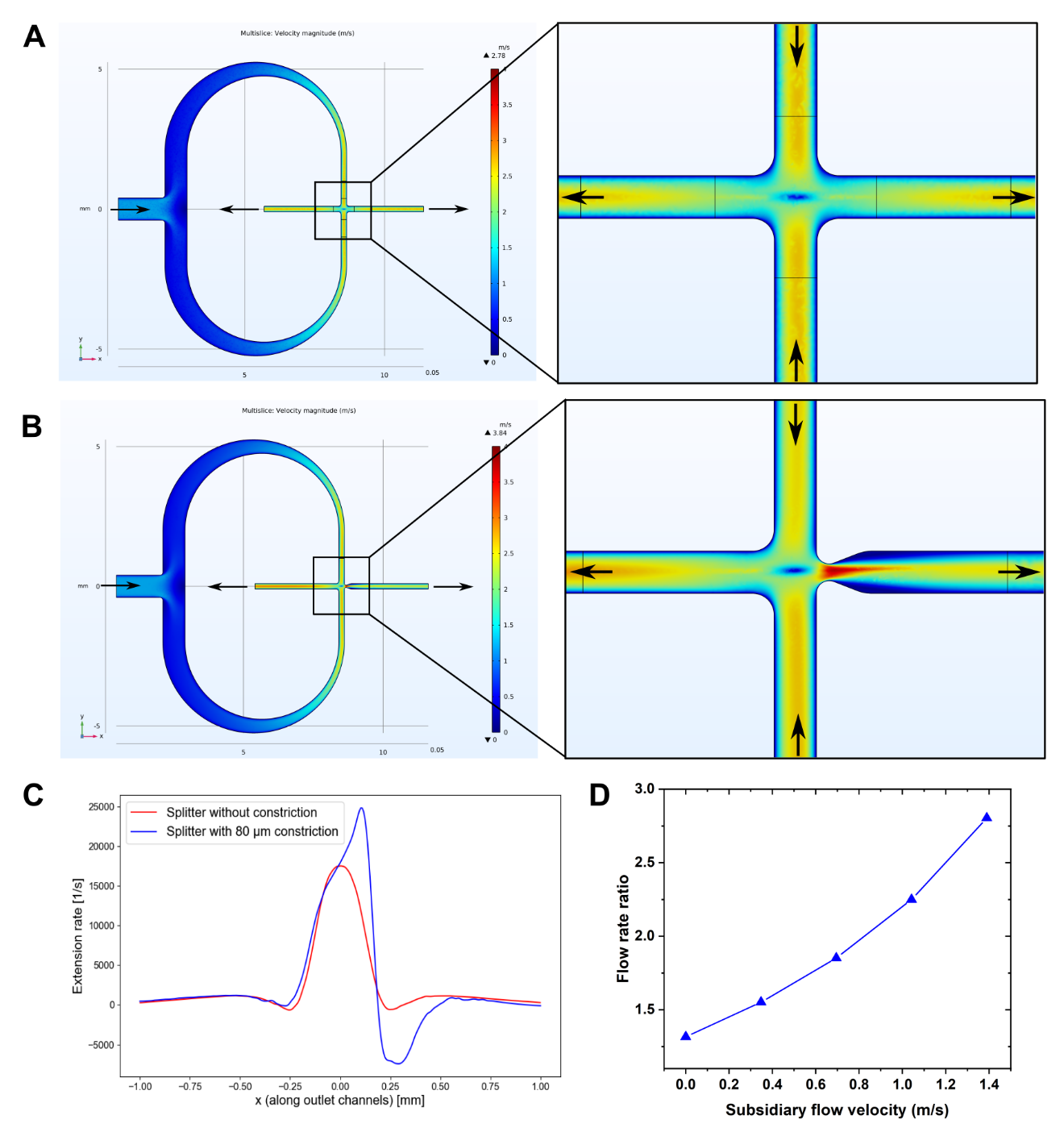

**Fig. S4. Numerical simulation of flow inside hydrodynamic splitter. A.** Numerical simulation of flow inside the symmetric hydrodynamic splitter shows a symmetric flow distribution at the cross junction. We used COMSOL Multiphysics 6.0 to run no-slip laminar flow simulations in 3D solving the Navier-Stokes constitutive equations. We used fine tetrahedral meshing (minimum element size 12.1 µm) with refinement near the boundary layers and at the cross junction. P2 + P2 discretization was used for solving the governing equations. **B.** Asymmetric flow is observed when a constriction is added at one of the outlet branches. The width of the channel at the constriction is 80 µm. We used a main flow rate of 200 mL/h in both simulations and the graphs were plotted using the same colorbar in both figures. Arrows denote the direction of flow in both simulations. **C.** The extension rate at the cross junction is plotted for the mid plane as a function of the horizontal coordinate x. The origin is located at the geometric center of the cross junction. The extension rate is symmetrically distributed around the center of the cross junction for the symmetric splitter without any constriction. Presence of a constriction increases the extension rate on the side where the constriction is present. The increase in extension rate by virtue of addition of a constriction is expected to increase the probability of splitting of cells at the cross junction. **D.** Numerical simulation of flow at the cross junction of the asymmetric hydrodynamic splitter shows an increasing ratio of flow rate on either side of the cross junction with subsidiary flow velocity. The flow rate ratio is defined as the ratio of the volumetric flow rate on the left side of the cross junction to the right side of the cross junction at a location to the left of the subsidiary flow channel. The main flow rate used in this simulation was 200 mL/h. Other simulation parameters were identical to those used previously.

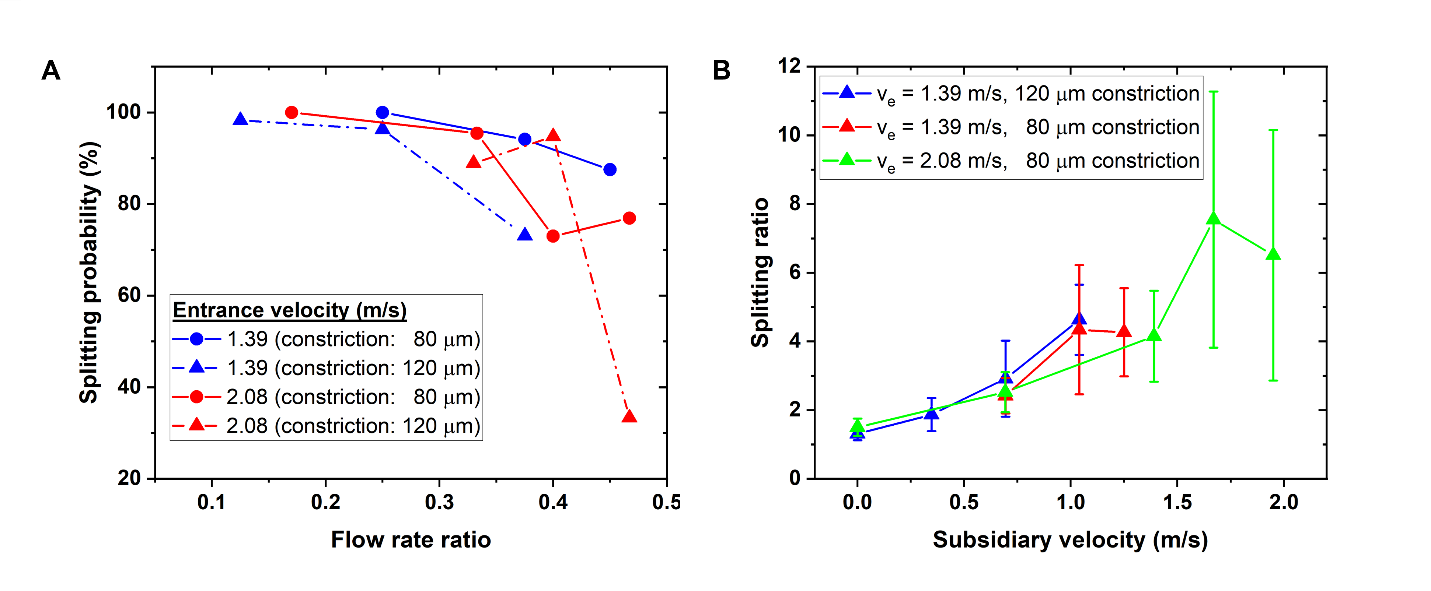

**Fig. S5. Effect of constriction in asymmetric hydrodynamic splitter. A.** Cell splitting probability is plotted as a function of the flow rate ratio for two entrance velocities and two constriction channel widths, where the flow rate ratio is defined as the ratio of the subsidiary flow rate to the main flow rate. The constriction channel width is defined as the width of the channel at the constriction. **B.** Splitting ratio as a function of the subsidiary velocity for different constriction channel widths and entrance velocities. The constriction channel width does not impact the splitting ratio significantly but allows us to operate the device at higher subsidiary flow velocities because of the higher splitting probability.

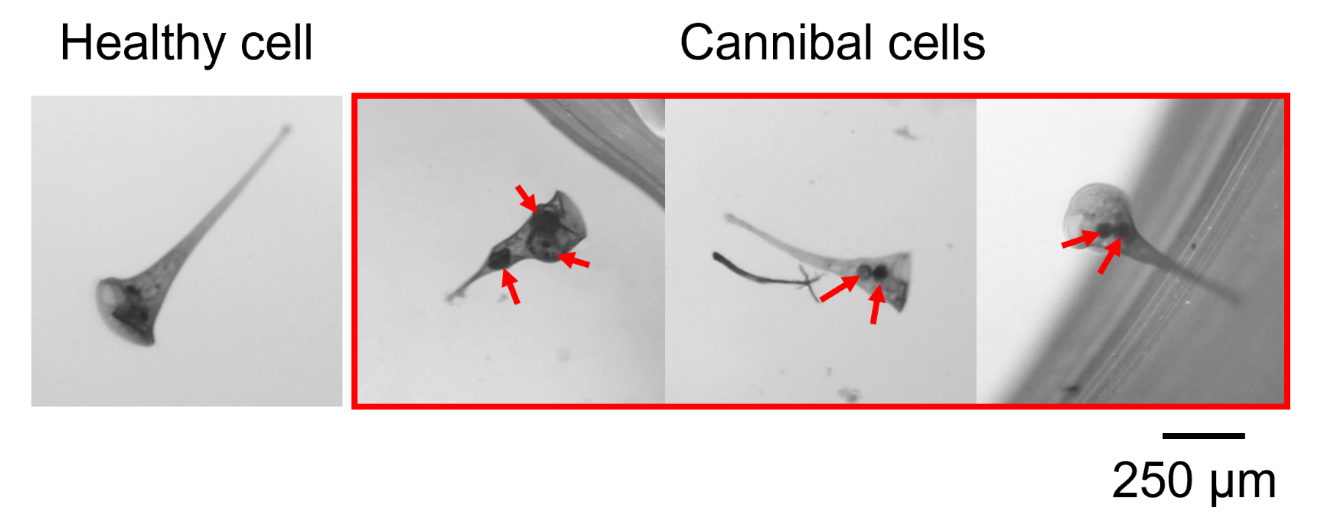

**Fig. S6. Cannibalism in *Stentors*.** The red box shows images of three cannibal *Stentor* cells with engulfed cell fragments (red arrows). A healthy *Stentor* cell is shown in the left panel for comparison.

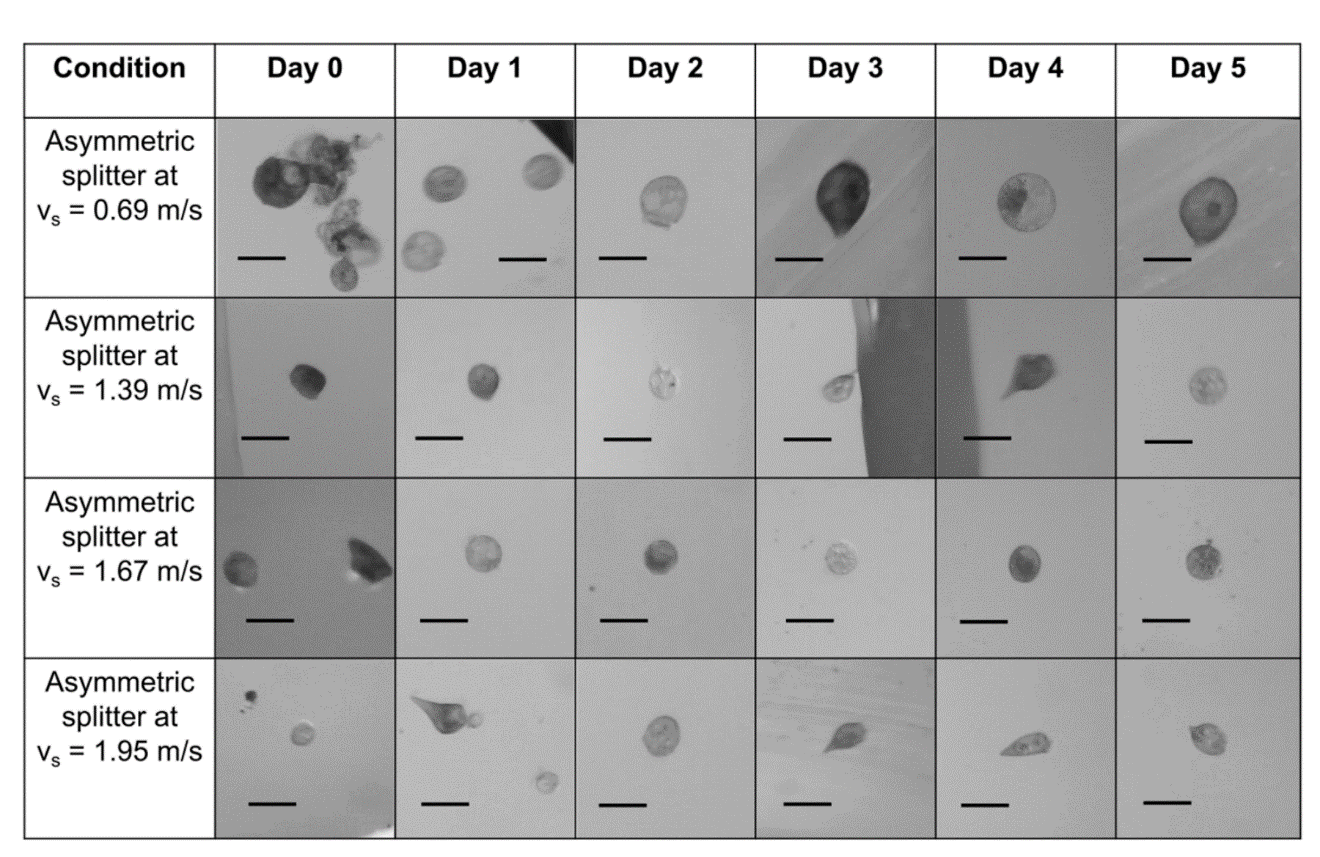

**Fig. S7.** Table showing representative images of unhealthy-looking but living cells (see definition in the main text) in the 5-day survival assay for different devices and flow conditions. The asymmetric splitter was operated at an entrance velocity of 2.08 m/s. For the guillotine and the asymmetric splitter without a subsidiary flow, all cell fragments appeared healthy were not included in this table (see Fig. 4F instead). The cell fragments obtained from the asymmetric splitter at v_s_ = 1.95 m/s are identical to the ones shown in Fig. 4F of the main text, because the fragments generated under this condition looked unhealthy in general and very few cells were viable. The scale bar in all images is 125 µm.

**Movie S1.** Movie showing a *Stentor* cell splitting at an entrance velocity of 2.08 m/s at the cross junction of a symmetric hydrodynamic splitter. The flapping of the tail end of each fragment post-splitting could be caused by formation of streamwise vortices downstream of the cross junction.

**Movie S2.** Movie showing one of the smallest live *Stentor* fragments generated from the splitting process. This fragment was formed after splitting using a symmetric hydrodynamic splitter at an entrance velocity of 2.08 m/s. The fragment is relatively transparent optically, has beating cilia and is membrane bound.
